# Activation of glucocorticoid receptor signaling inhibits KSHV-induced inflammation and tumorigenesis

**DOI:** 10.1101/2023.11.10.566578

**Authors:** Luping Chen, Ling Ding, Xian Wang, Yufei Huang, Shou-Jiang Gao

## Abstract

Hyperinflammation is the hallmark of Kaposi’s sarcoma (KS), the most common cancer in AIDS patients caused by Kaposi’s sarcoma-associated herpesvirus (KSHV) infection. However, the role and mechanism of induction of inflammation in KS remain unclear. In a screening for inhibitors of KSHV-induced oncogenesis, over half of the identified candidates were anti-inflammatory agents including dexamethasone functions by activating glucocorticoid receptor (GR) signaling. Here, we examined the mechanism mediating KSHV-induced inflammation. We found that numerous inflammatory pathways were activated in KSHV-transformed cells. Particularly, interleukin-1 alpha (IL-1α) and IL-1 receptor antagonist (IL-1Ra) from the IL-1 family were the most induced and suppressed cytokines, respectively. We found that KSHV miRNAs mediated IL-1α induction while both miRNAs and vFLIP mediated IL-1Ra suppression. Furthermore, GR signaling was inhibited in KSHV-transformed cells, which was mediated by vFLIP and vCyclin. Dexamethasone treatment activated GR signaling, and inhibited cell proliferation and colony formation in soft agar of KSHV-transformed cells but had a minimal effect on matched primary cells. Consequently, dexamethasone suppressed the initiation and growth of KSHV-induced tumors in mice. Mechanistically, dexamethasone suppressed IL-1α but induced IL-1Ra expression. Treatment with recombinant IL-1α protein rescued the inhibitory effect of dexamethasone while overexpression of IL-1Ra caused a weak growth inhibition of KSHV-transformed cells. Furthermore, dexamethasone induced IκBα expression resulting in inhibition of NF-κB pathway and IL-1α expression. These results reveal an important role of IL-1 pathway in KSHV-induced inflammation and oncogenesis, which can be inhibited by dexamethasone-activated GR signaling, and identify IL-1-mediated inflammation as a potential therapeutic target for KSHV-induced malignancies.

**Importance:** Kaposi’s sarcoma (KS) is the most common cancer in HIV-infected patients caused by Kaposi’s sarcoma-associated herpesvirus (KSHV) infection. Hyperinflammation is the hallmark of KS. In this study, we have shown that KSHV mediates hyperinflammation by inducing IL-1α and suppressing IL-1Ra. Mechanistically, KSHV miRNAs and vFLIP induce hyperinflammation by activating the NF-κB pathway. A common anti-inflammatory agent dexamethasone blocks KSHV-induced hyperinflammation and tumorigenesis by activating glucocorticoid receptor signaling to suppress IL-1α and induce IL-1Ra. This work has identified IL-1-mediated inflammation as a potential therapeutic target and dexamethasone as a potential therapeutic agent for KSHV-induced malignancies.

## Introduction

Kaposi’s sarcoma (KS), the most common cancer in AIDS patients caused by infection of Kaposi’s sarcoma-associated herpesvirus (KSHV), is a vascular spindle cell cancer (1). Although the incidence of KS in Western countries has declined in the era of anti-retroviral therapy, over half of AIDS-related KS (AIDS-KS) patients cannot achieve total remission (2, 3). In sub-Saharan Africa, KS remains as the predominant cancer in both immunocompetent and immunocompromised individuals (1). Anti-cancer therapies are often ineffective against KS, and there is no vaccine or anti-KSHV therapy that can prevent or eliminate persistent KSHV infection (3-5). As a result, KS continues to be an urgent cancer, causing significant mortality and morbidity in the affected populations worldwide (3, 6). In addition to KS, KSHV is etiologically associated with primary effusion lymphoma (PEL), a subset of multicentric Castleman’s disease (MCD), and KSHV inflammatory cytokine syndrome (KICS) (2, 7).

Numerous cofactors have been implicated in KSHV-associated malignancies. KSHV mostly causes cancers in patients with suppressed immunity, particularly in AIDS-related KS (2, 5). HIV products such as Tat, Vpr and Nef proteins also regulate KSHV replication and promote the proliferation of KS tumor cells (8-12). Iron has been implicated in endemic KS in sub-Saharan Africa (13, 14). Despite the roles of different cofactors in KS, all clinical forms of KS are characterized by uncontrolled hyperinflammation manifesting as vast immune cell infiltration, leaky vascular structures, and abundant inflammatory cytokines (15, 16). Nevertheless, the role of inflammation and the mechanism mediating inflammation in KS remain unclear.

Viral and bacterial coinfections, metabolic disorders, immunosuppression and abnormal activation of immune response in HIV-infected subjects contribute to KS hyperinflammation (17). In particular, oral bacterial infections induce inflammation and enhance the progression of oral KS (18-21).

KSHV infection also induces inflammation (2, 7). Similar to other herpesviruses, KSHV has a bi-phasic viral life cycle: latency and lytic replication. During latency, KSHV has restricted expression of latent genes/products including latency-associated nuclear antigen (LANA), viral cyclin (vCyclin), viral FLICE inhibitory protein (vFLIP), and a cluster of 12 viral precursor microRNAs (miRNAs) (22, 23). During lytic replication, KSHV expresses most of the lytic genes and produces infectious virions. In the early stage of AIDS-KS, most tumor cells are latently infected by KSHV; however, a small number of them also undergo spontaneous lytic replication, producing viral cytokines/chemokines and infectious virions (7, 23). In fact, high levels of viral interleukin-6 (vIL-6) and cellular cytokines can be detected during KSHV lytic replication (24-27). The infectious virions infect new cells and induce proinflammatory and proangiogenic cytokines, such as matrix metalloproteinases (MMPs), IL-6, angiopoietins, and IL-8 (28-31). Indeed, clinical studies have reported that KS progression is correlated with KSHV loads and lytic antibody titers (32-35). Thus, the expression of viral lytic genes promotes cell proliferation and inflammation, which is essential for the initiation and progression of the early stage of KS tumors (7, 23). Nevertheless, most tumor cells are latently infected by KSHV and there is almost no viral lytic cell in advanced stage of KS (1, 2, 7, 23), suggesting the essential role of latent infection in KS development. In fact, KSHV latent products are required for maintaining viral latency, promoting host cell proliferation and survival, and inducing proinflammatory cytokines (7, 22, 23).

Despite being a pathological hallmark, the role of inflammation in KS development remains unclear. Inflammation and oxidative stress can induce and promote KSHV lytic replication, thus likely contributing to the development of early stage of KS (36, 37). As KSHV-encoded vFLIP and miRNAs can promote cell survival (7, 23), KSHV-infected cells are refractory to inflammation and oxidative stress, which likely, on the contrary, promote cell proliferation, malignant transformation and tumorigenesis (17, 38-40).

In an effort to identify inhibitors of KSHV-induced oncogenesis, we have previously performed a high-throughput screening (HTS) of 3,731 characterized compounds with KSHV-transformed rat primary mesenchymal precursor cells (KMM) and matched primary cells (MM) (41, 42). Surprisingly, over half of the identified inhibitors are anti-inflammatory, including dexamethasone. Dexamethasone is a corticosteroid widely used in many conditions for its anti-inflammatory and immunosuppressive properties (43). It is an agonist of the glucocorticoid receptor (GR), a transcription factor regulating numerous cellular signaling pathways (43).

In this study, we examined the role of inflammation in KSHV-induced cellular transformation and tumorigenesis, and determined the mechanism of KSHV induction of inflammation. We also examined the effect of dexamethasone on KSHV-transformed cells and determined its therapeutic potential. We found that dexamethasone could effectively inhibit the proliferation and cellular transformation of KMM cells with minimal effect on MM cells. Dexamethasone also inhibited the initiation and growth of KS-like tumors. RNA-seq revealed that KSHV infection induced inflammatory cytokines such as IL-1α and suppresses anti-inflammatory cytokines such as IL-1Ra. Dexamethasone inhibited KSHV induction of IL-1α and reversed KSHV suppression of IL-1Ra by activating the GR signaling pathway. Furthermore, dexamethasone blocked the nuclear factor kappa B (NF-κB) signaling activated by KSHV infection through GR-transactivation of IκBα. Together, these results revealed the important roles of proinflammatory IL-1 and NF-κB signaling in KSHV-induced oncogenesis, and that activation of the anti-inflammatory GR signaling with dexamethasone and likely other anti-inflammatory drugs could be a promising therapeutic approach for KSHV-induced malignancies.

## Results

### Inflammatory pathways are activated in KSHV-transformed cells

To determine whether the inflammatory pathways were activated in KSHV-transformed cells, we performed mRNA-seq and examined the differentially expressed genes (DEGs) in KSHV-transformed cells. Using the cutoffs of fold change of 2.5 and FDR of 0.05, we observed 1,425 downregulated DEGs and 1,551 upregulated DEGs in KMM cells compared to MM cells (Fig. 1A and Table S1), which manifested in clusters in the heatmap (Fig. 1B). Numerous enriched clusters were related to immune and inflammatory response. Gene ontology biological process (GOBP) pathway analysis identified the enriched pathways of KSHV-transformed cells, including wound healing, positive regulations of interleukin-8 (IL-8) production, positive regulation of phosphatidylinositol 3 kinase (PI3K) signaling, angiogenesis, response to virus, positive regulation of mitogen-activated protein kinase (MAPK) cascade, cellular response to IL-1, response to hypoxia, and inflammatory response (Fig. 1C and Table S2). Taken together, these results showed that multiple inflammatory pathways were activated in KSHV-transformed cells.

**FIG 1.**
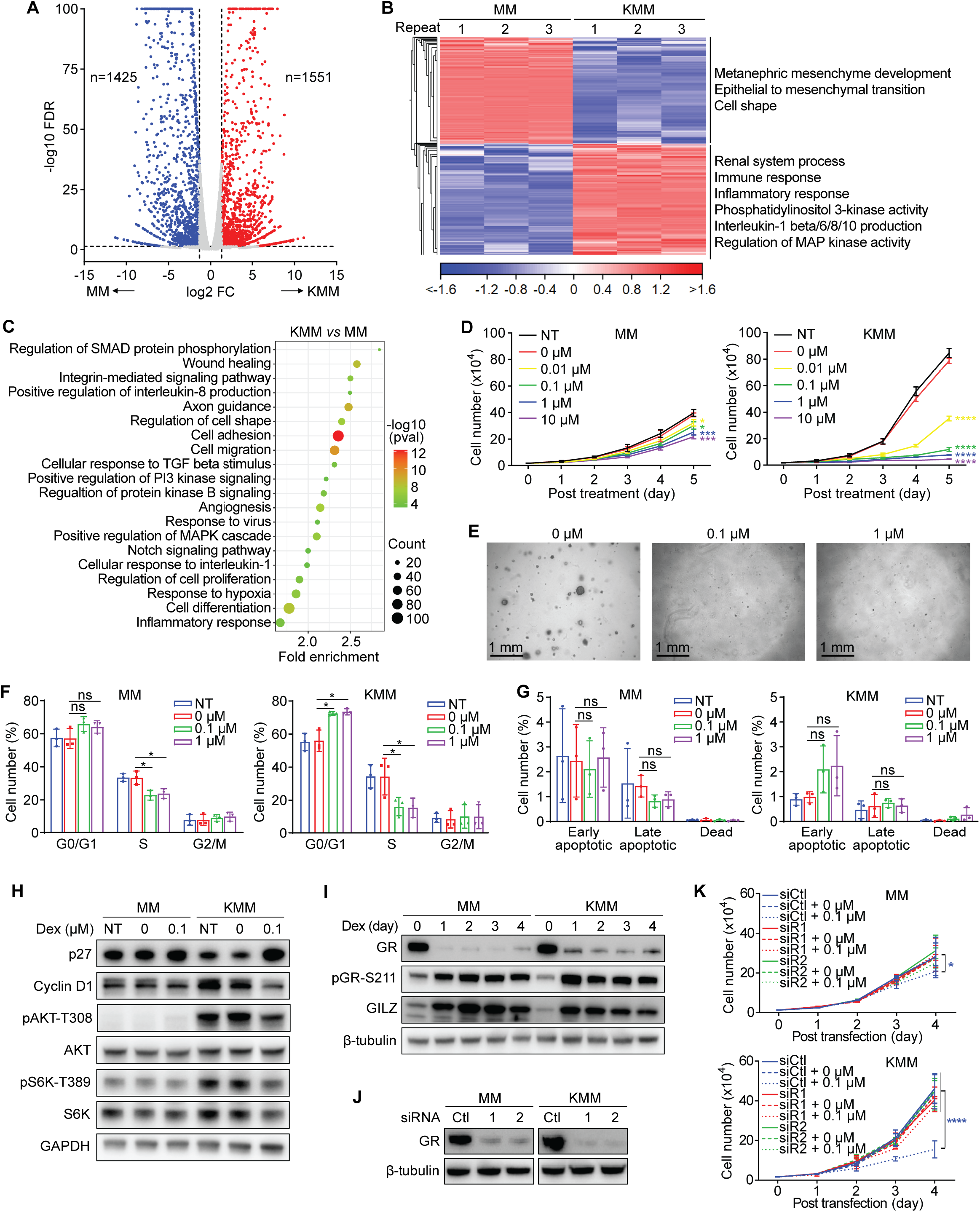
Dexamethasone suppresses inflammatory pathways, and inhibits KSHV-induced cell proliferation and cellular transformation. (A) Volcano plot of differentially expressed genes (DEGs) between KSHV-transformed KMM cells and matched primary MM cells. Cutoffs: Fold change > 2.5, FDR < 0.05, P < 0.05. (B) Heatmap of DEGs between MM and KMM cells. (C) Bubble chart of enriched GOBP pathways. (D) Cell proliferation curves of MM and KMM cells treated with 0, 0.01, 0.1, 1 or 10 µM dexamethasone dissolved in 0.1% ethanol. (E) Representative pictures of soft agar assay of KMM cells treated with 0, 0.1 or 1 µM dexamethasone for 15 days. Scare bar: 1 mm. (F and G) Analysis of cell cycle (F) and apoptosis (G) of MM and KMM cells treated with 0, 0.1 or 1 µM dexamethasone for 2 days. (H) Immunoblots of p27, cyclin D1, pAKT-T308, AKT, pS6K-T389 and S6K in MM and KMM cells treated with 0 or 0.1 µM dexamethasone for 2 days. (I) Immunoblots of GR, pGR-S211 and GILZ in MM and KMM cells treated with 0 or 0.1 µM dexamethasone. (J) Immunoblot of GR following siRNA knockdown in MM and KMM cells. (K) Cell proliferation curves of MM and KMM cells following GR knockdown treated with 0 or 0.1 µM dexamethasone. Experiments were independently repeated three times, and results are presented as mean ± s.d. from the three experiments (D, F, G and K).

### Dexamethasone inhibits the proliferation and cellular transformation of KSHV-transformed cells

Next, we determined whether suppression of inflammation with dexamethasone could inhibit KSHV-induced cellular transformation. Dexamethasone is a classic corticosteroid medication commonly used to relieve inflammation and treat a variety of inflammatory conditions such as arthritis, severe allergies, skin diseases, colitis, asthma, as well as COVID-19 (44-46). Treatment with dexamethasone significantly inhibited the proliferation of KMM cells in a concentration-dependent fashion but only had a marginal effect on MM cells (Fig. 1D). Dexamethasone at 0.1 µM was sufficient to almost completely inhibit the proliferation and abolished colony formation in soft agar of KMM cells (Fig. 1E). Dexamethasone induced cell cycle arrest in KMM cells by increasing G0/G1-phase cells from 55-57% to 72-74% and decreasing S-phase cells from 34-35% to 16-17% at both 0.1 and 1 µM concentrations (Fig. 1F). Minor effects on cell cycle progression in MM cells were observed under the same conditions. In contrast, dexamethasone had minimal effect on the survival of both MM and KMM cells (Fig. 1G). We also did not detect any increases of cleaved PARP, and cleaved caspases 3 and 7 in both MM and KMM cells following dexamethasone treatment (results not shown).

Consistent with the faster proliferation rate, KMM cells have a lower level of p27 and a higher level of cyclin D1 than MM cells have (47). Dexamethasone at 0.1 µM was sufficient to significantly increase the level of p27 and suppress the level of cyclin D1 in KMM cells but had no noticeable effect on MM cells (Fig. 1H). Among the mitogenic and oncogenic pathways activated by KSHV infection, the AKT/mTOR pathway regulates the cell cycle progression (48). As expected, treatment with 0.1 µM dexamethasone significantly reduced the levels of pAKT-T308 and pS6K-T389 in KMM cells (Fig. 1H).

### GR signaling is required for dexamethasone inhibition of KSHV-induced cell proliferation

Dexamethasone binds GR to activate its downstream signaling pathways (43, 49). Thus, we determined whether GR signaling was functional in MM and KMM cells. Dexamethasone treatment increased the levels of GR phosphorylation at S211 (pGR-S211) and downstream GR transcriptional target glucocorticoid-induced leucine zipper (GILZ) in MM and KMM cells (Fig. 1I), indicating that GR signaling was functional in both types of cells. As expected, dexamethasone also reduced the total GR protein level in both MM and KMM cells, which was due to proteasome-mediated degradation following signaling activation by ligand binding to the receptor (50). GR knockdown by siRNAs completely abolished the inhibitory effect of dexamethasone on the proliferation of KMM cells (Fig. 1J and K). Thus, GR signaling was functional and necessary for dexamethasone suppression of KSHV-induced cell proliferation.

### Dexamethasone suppresses the initiation and growth of KSHV-induced tumors

We further investigated whether dexamethasone could inhibit KSHV-induced tumorigenesis with the KMM model. The experiment was performed in two stages. In stage I, we examined the effect of dexamethasone on tumor incidence and growth (Fig. 2A). Nude mice were subcutaneously inoculated with KMM cells on both flanks to induce KS-like tumors. We observed palpable tumors after 17 days. Half of the mice (n=16) then received the dexamethasone administration (Dex) 5 days per week for 5 weeks, while the other half (n=16) was treated with solvent (vehicle). When using a volume of 100 mm^3^ as the threshold for tumor incidence, mice treated with dexamethasone showed a lower tumor incidence throughout the experiment compared to the vehicle group (Fig. 2B). The vehicle group reached a 50% incidence rate on day 23 while it took 38 days for the dexamethasone group to reach the same rate (P < 0.01 for trend). At the end of stage I experiment at day 53, all the inoculated sites (32 out of 32) developed tumors in the vehicle group while only 25 out of 32 (78.13%) of the inoculated sites developed tumors in the dexamethasone group. Dexamethasone also significantly inhibited the growth rate of tumors (Fig. 2C). At the end of the experiment, the average tumor size was significantly smaller in the dexamethasone group than that in the vehicle group (128.11 *vs* 203.45 mm^3^, P < 0.0001). Meanwhile, dexamethasone treatment caused a slight weight loss compared to the vehicle group (Fig. S1A).

**FIG 2.**
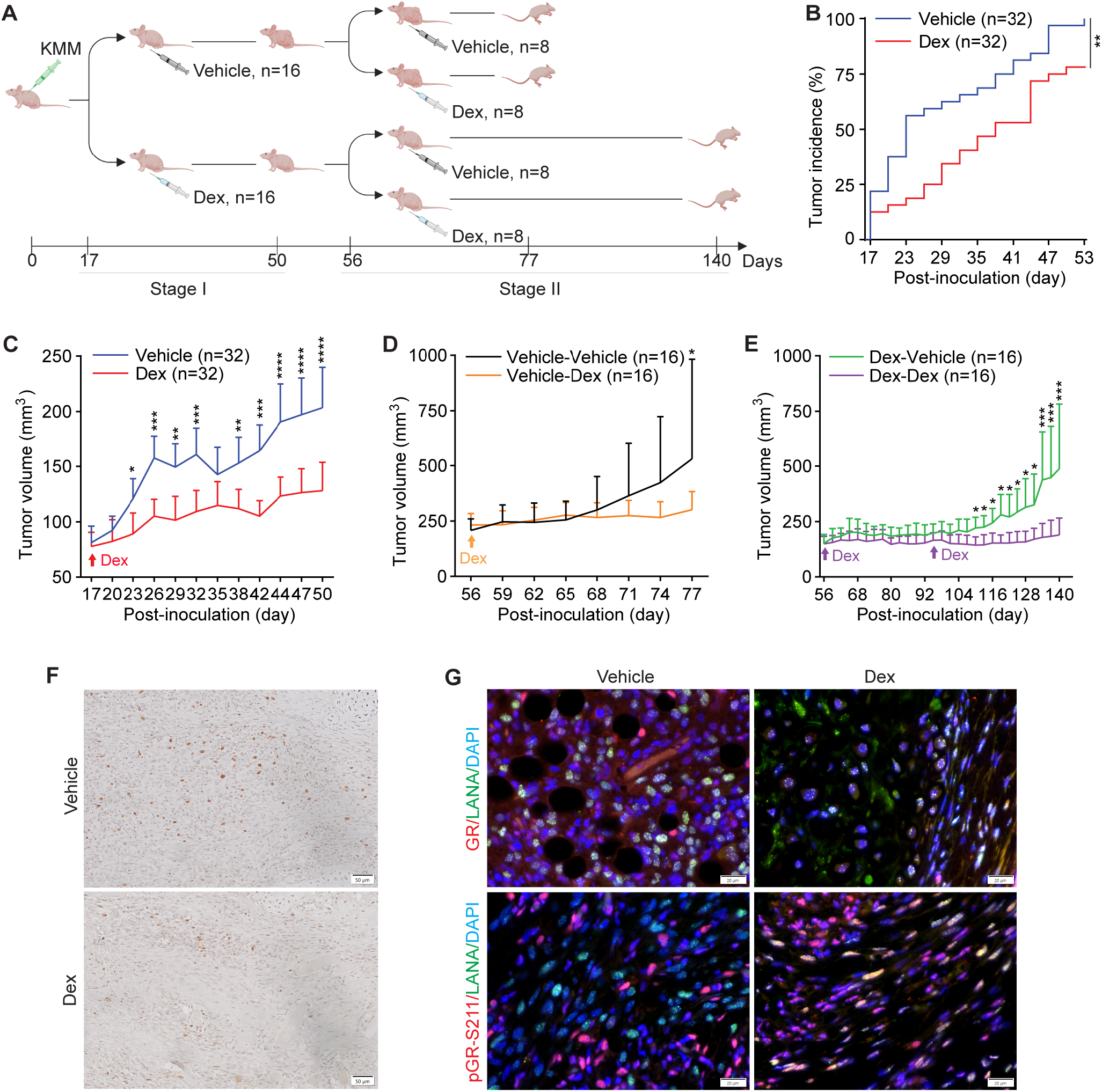
Dexamethasone suppresses tumor initiation and growth of KS-like tumors. (A) Schematic illustration of experimental design. Stage I was designed to examine the effect of dexamethasone on tumor initiation and growth while Stage II was designed to examine the effect of dexamethasone on grown tumors and maintenance of dexamethasone-treated tumors. Note that mice were inoculated on both flanks to generate 2 tumors per mouse. (B) Tumor incidence over time in vehicle- and dexamethasone-treated groups during Stage I experiment. Each group contained 32 tumors from 16 mice. Log rank test was used for the trend analysis. (C) Tumor growth of vehicle- and dexamethasone-treated groups during Stage I experiment. Each group contained 32 tumors from 16 mice. (D) Tumor growth of vehicle-treated mice from Stage I experiment and further treated with either vehicle or dexamethasone in Stage II experiment. Each group contained 16 tumors from 8 mice. (E) Tumor growth of dexamethasone-treated mice from Stage I experiment and further treated with either vehicle or dexamethasone in Stage II experiment. Each group contained 16 tumors from 8 mice. (F) Ki67 staining of the KS-like tumors. Scare bar: 50 µM. (G) Dual immunofluorescence staining of LANA and GR, or LANA and pGR-S211 of the KS-like tumors. Scare bar: 20 µM. Student’s t test was used for the analysis and results are presented as mean + 95% CI (C-E).

In stage II, we examined whether dexamethasone could regress grown tumors or maintain its inhibitory effect on tumor growth after stopping the treatment (Fig. 2A). We used the vehicle group from stage I to examine whether dexamethasone could regress grown tumors. Mice from this group were evenly divided into two subgroups, and one subgroup was treated with dexamethasone (Vehicle-Dex, n=8) while the second subgroup remained untreated (Vehicle-Vehicle, n=8). While the tumors in the Vehicle-Vehicle subgroup continued the aggressive growth, those in the Vehicle-Dex group stopped to grow (Fig. 2D). At the end of the experiment at day 77, the average tumor size was significantly smaller in the Vehicle-Dex group than that in the Vehicle-Vehicle group (510 *vs* 255 mm^3^, P < 0.05). Nevertheless, dexamethasone failed to regress the grown tumors.

We used the dexamethasone group from stage I to examine whether the inhibitory effect of dexamethasone on tumor growth could be maintained after stopping the treatment. Mice from this group were divided into two subgroups, and one subgroup remained treated with dexamethasone (Dex-Dex, n=8) while the second subgroup was no longer treated (Dex-Vehicle, n=8). While tumors in the Dex-Vehicle group were maintained in the same volumes for more than 7 weeks, they ultimately started to grow (Fig. 2E). In contrast, tumors in the Dex-Dex group had minimal growth during the entire experiment (Fig. 2E). Similar to the results of stage I, a small loss of mouse weight was observed with dexamethasone administration (Fig. S1B and C).

Together, these results indicated that dexamethasone treatment reduced tumor incidence, inhibited tumor growth of both small and grown tumors, and maintained its inhibitory effect on tumor growth for more than 7 weeks after stopping the treatment. These results were in agreement with dexamethasone’s inhibitory effect on cell cycle progression but not apoptosis (Fig. 1F and G). Indeed, examination of cell proliferation marker Ki67 revealed that dexamethasone-treated tumors had much fewer Ki67 positive tumor cells than those of vehicle-treated tumors (Fig. 2F). Immunofluorescence staining revealed that GR protein level was substantially lower in dexamethasone-treated than vehicle-treated tumors (Fig. 2G), which was consistent with the downregulation of GR protein in KMM cells following dexamethasone treatment (Fig. 1I). Interestingly, there was almost no detectable signal of pGR-S211 in the LANA-positive cells albeit strong signals in LANA-negative cells in vehicle-treated tumors (Fig. 2G), suggesting the presence of a negative regulatory mechanism of GR signaling in the basal level in the tumors without any dexamethasone stimulation. Thus, we reexamined GR signaling in MM and KMM cells, and indeed observed a lower pGR-S211 level in KMM than MM cells without dexamethasone treatment though both cell lines responded well to dexamethasone treatment (Fig. 1I). Consistent with these results, we detected robust pGR-S211 levels in almost all cells in the tumors under dexamethasone treatment, thus confirming the responsiveness of tumor cells to dexamethasone (Fig. 2G).

### Dexamethasone inhibits inflammatory response pathways in KSHV-transformed cells

In order to delineate the mechanism of dexamethasone inhibition of proliferation of KSHV-transformed cells, we performed mRNA-seq on both MM and KMM cells treated with 0.1 µM dexamethasone for 0 (untreated), 2 and 4 days. Principal component analysis (PCA) showed consistent results among three biological replicates of each group (Fig. 3A). As expected, MM and KMM cells were separated into two clusters. Cells treated for 2 and 4 days generally manifested more similar profiles than those of untreated cells. Thus, we generated a union of day 2 and 4 samples for gene expression analysis and compared them with day 0 samples. We identified 735 downregulated DEGs and 1,079 upregulated DEGs in KMM cells, and 692 downregulated DEGs and 1,393 upregulated DEGs in MM cells, respectively (Fig. 3B; Supplementary Table S3 and S4). There were 257 downregulated genes and 534 upregulated genes that were shared between MM (37.14% and 38.33%) and KMM (34.97% and 49.49%) cells, respectively (Fig. 3C), suggesting distinct responses of these two cell types to dexamethasone. The heatmaps confirmed the reproducibility by showing that three repeats at different time points in both MM and KMM cells clustered together (Fig. 3D). As expected, similar clustering patterns were observed between day 2 and 4 following dexamethasone treatment in both MM and KMM cells. Many altered genes were related to inflammatory response and GR signaling, particularly in KMM cells (Fig. 3D). GOBP analysis confirmed that the response to glucocorticoid pathway was enriched in both MM and KMM cells following dexamethasone treatment (Fig. 3E; Supplementary Table S5 and S6). Importantly, many inflammatory pathways were enriched in the KMM cells, including cellular responses to IL-1, wound healing, cellular response to tumor necrosis factor (TNF), cellular response to interferon-γ (IFN-γ), inflammatory response, and response to lipopolysaccharide (LPS), etc (Fig. 3E). In particular, the GSEA hallmark of inflammatory response and the KEGG cell cycle pathway were enriched in KMM cells but only had marginal effect on MM cells following dexamethasone treatment (Fig. 3F). Thus, dexamethasone effectively targeted inflammatory responses and cell cycle pathways in KMM cells while the effects on MM cells were minimal.

**FIG 3.**
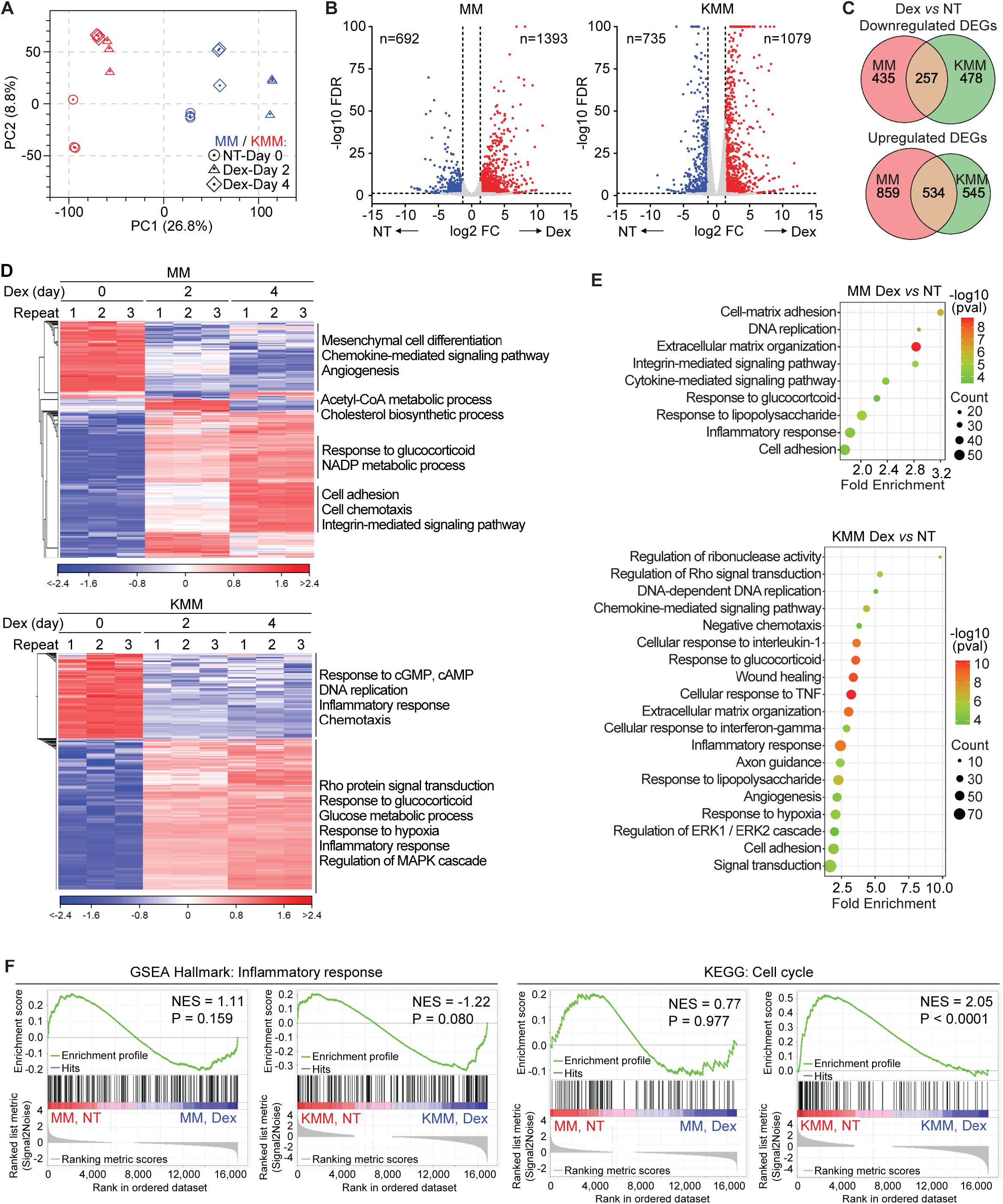
Dexamethasone suppresses inflammatory pathways of KSHV-transformed cells. (A) PCA plot of mRNA-seq results of MM and KMM cells treated with 0.1 µM dexamethasone for 0, 2 and 4 days. (B) Volcano plots of differentially expressed genes (DEGs) in MM and KMM cells treated with dexamethasone described in (A). Cutoffs: Fold change > 2.5, FDR < 0.05, P < 0.05. (C) Venn diagrams showing overlaps of upregulated and downregulated DEGs of MM and KMM cells treated with dexamethasone. (D) Heatmaps of DEGs of MM and KMM cells treated with dexamethasone. (E) Bubble chart of enriched GOBP pathways in MM and KMM cells treated with dexamethasone. (F) Enrichment plots of inflammatory response (GSEA hallmark) and cell cycle (KEGG) of MM and KMM cells treated with dexamethasone.

### Dexamethasone inhibits KSHV induction of IL-1α and releases KSHV suppression of IL-1Ra

IL1 signaling was one of the enriched inflammatory pathways affected in KMM cells following treatment with dexamethasone (Fig. 3E). IL-1α and IL-1Ra were the two top genes in this pathway significantly regulated by dexamethasone (Table S5 and S6). IL-1α is a proinflammatory cytokine whose function depends on its binding to the IL-1 receptor, whereas IL-1Ra is the natural antagonist of IL-1α that binds to the same receptor but functions to inhibit IL-1 signaling (51). The mRNA-seq results showed that IL-1α transcript was upregulated 5-fold in KMM cells compared to MM cells (Fig. 4A), which was confirmed by RT-qPCR (Fig. 4B). An ELISA assay confirmed the upregulation of IL-1α protein in KMM cells compared to MM cells (Fig. 4C). However, treatment with dexamethasone almost completely abolished KSHV induction of IL-1α transcript and protein (Fig. 4A-C). These results indicated that KSHV infection induced the expression of proinflammatory IL-1α, which was suppressed by dexamethasone.

**FIG 4.**
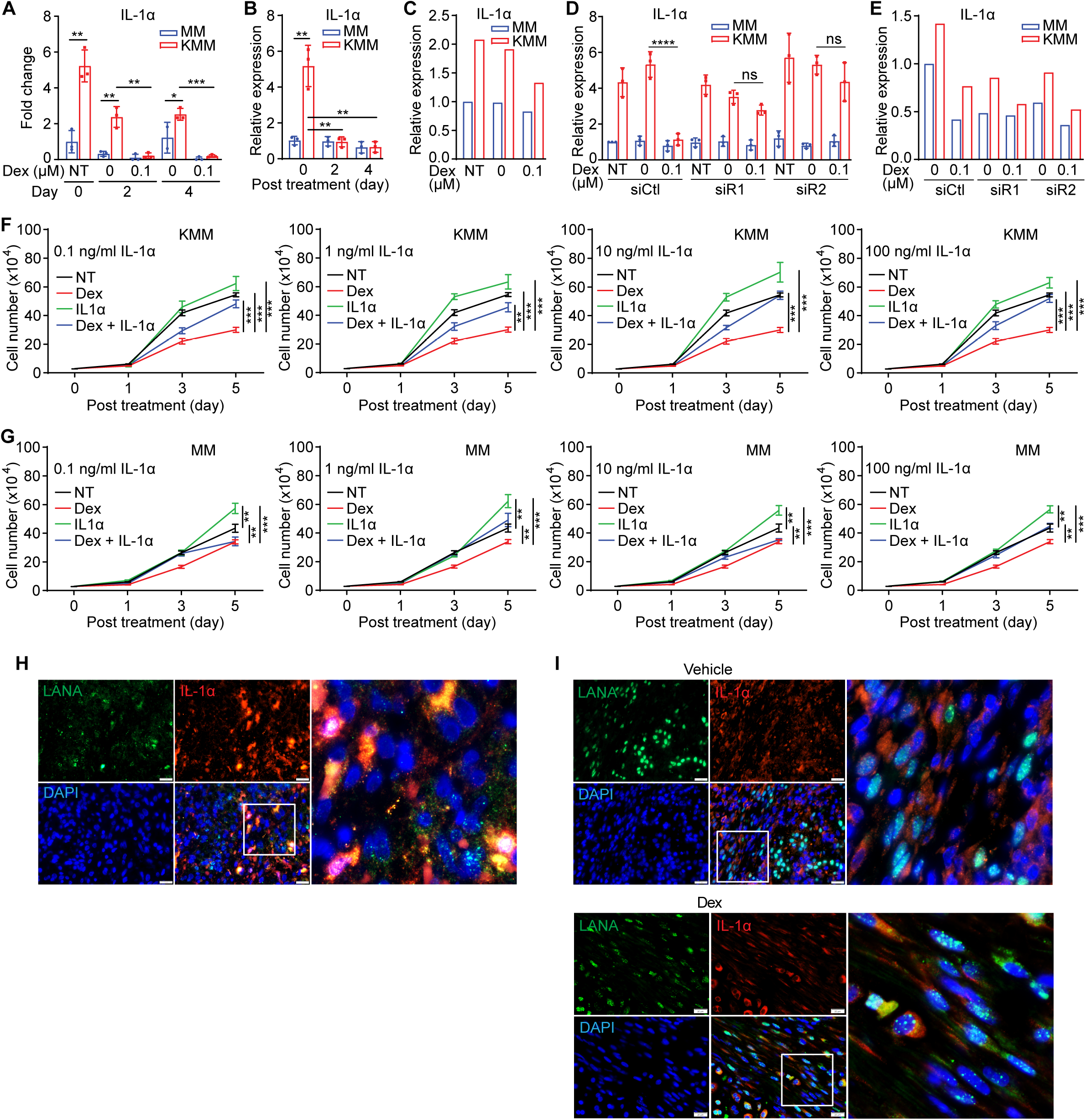
IL-1α mediates dexamethasone suppression of cell proliferation in KSHV-transformed cells. (A and B) IL-1α mRNA expression in MM and KMM cells treated with 0 or 0.1 µM dexamethasone measured by RNA-seq (A) and RT-qPCR (B). (C) IL-1α protein level in MM and KMM cells treated with 0 or 0.1 µM dexamethasone for 2 days detected by ELISA. (D) IL-1α mRNA expression in MM and KMM cells following knockdown of GR treated with 0 or 0.1 µM dexamethasone for 2 days measured by RT-qPCR. Cells were treated with dexamethasone one day after siRNA transfection. (E) IL-1α protein level in MM and KMM cells following knockdown of GR treated with 0 or 0.1 µM dexamethasone for 2 days detected by ELISA. (F and G) Cell proliferation curves of KMM (F) and MM (G) cells treated with 0 or 0.1 µM dexamethasone, and 0.1, 1, 10 or 100 ng/ml recombinant IL-1α protein. (H and I) Dual immunofluorescence staining of LANA and IL-1α in human KS tumors (H), or vehicle- and dexamethasone-treated mouse KS-like tumors (I). Scale bar: 20 µM. Student’s t test was used for the analysis and results from 3 independent repeats are presented as mean ± s.d. (B, D, F and G).

Next, we examined whether GR signaling was required for dexamethasone suppression of KSHV induction of IL-1α. Knockdown of GR effectively prevented dexamethasone suppression of IL-1α transcript and protein in KMM cells, albeit it did not affect the basal IL-1α expression in both MM and KMM cells (Fig. 4D and E).

To further determine whether IL-1α suppression was required for dexamethasone inhibition of proliferation of KSHV-transformed cells, we cultured the cells with dexamethasone or recombinant IL-1α protein alone, or both. IL-1α protein alone promoted the proliferation of KMM cells in a concentration-dependent manner, and reversed the inhibitory effect caused by dexamethasone (Fig. 4F). Similar results were also observed in MM cells but the effect was less obvious (Fig. 4G).

Moreover, we examined human KS lesions for IL-1α expression. LANA staining was used to identify KSHV-infected cells. A high level of IL-1α protein expression was observed in human KS lesions with (95.14±6.11)% LANA-positive cells expressing IL-1a (Fig. 4H). Examination of KS-like tumors from the KMM mouse model also revealed a high expression level of IL-1α protein in tumors from the vehicle group with (98.52±1.22)% LANA-positive cells expressing IL-1a (Fig. 4I). However, dexamethasone treatment reduced the overall IL-1α protein level as expected (Fig. 4I).

In contrast to IL-1α, the mRNA-seq results showed that the expression of IL-1Ra transcripts was downregulated 25-fold in KMM cells compared to MM cells (Fig. 5A), which was confirmed by RT-qPCR (Fig. 5B). Treatment with dexamethasone increased the expression levels of IL-1Ra transcripts in both MM and KMM cells (Fig. 5A and B), which was almost completely prevented following GR knockdown (Fig. 5C). Therefore, dexamethasone induction of IL-1Ra depended on the functional GR signaling in both types of cells.

**FIG 5.**
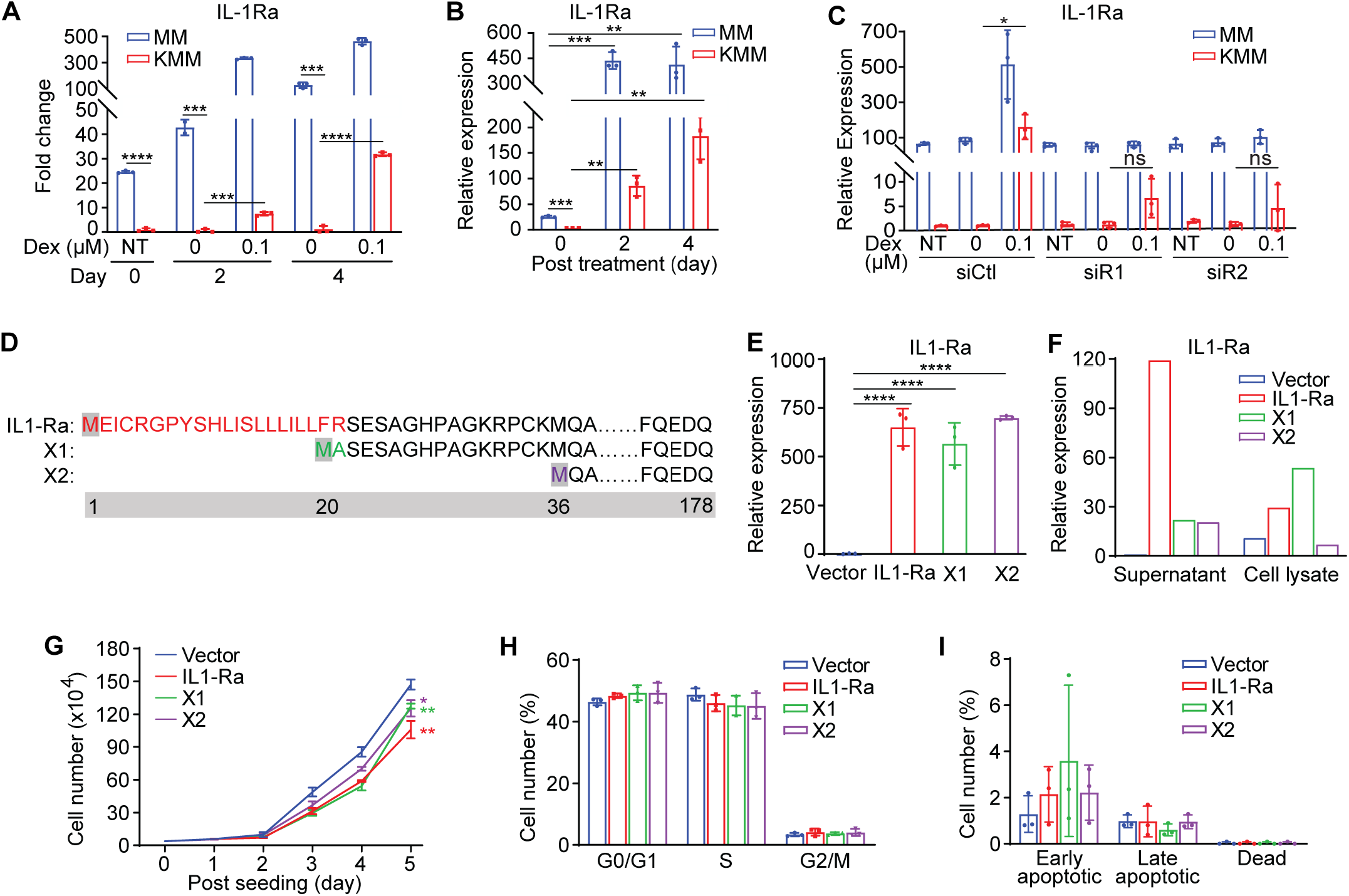
Induction of IL-1Ra marginally mediates dexamethasone suppression of cell proliferation in KSHV-transformed cells. (A and B) IL-1Ra mRNA expression in MM and KMM cells treated with 0 or 0.1 µM dexamethasone for 2 days measured by RNA-seq (A) and RT-qPCR (B). (C) IL-1Ra mRNA expression in MM and KMM cells following knockdown of GR treated with 0 or 0.1 µM dexamethasone measured by RT-qPCR. Cells were treated with 0.1 µM dexamethasone one day after siRNA transfection. (D) Schematic illustration of the differences between rat natural IL-1Ra protein, and predicted variants X1 and X2. (E and F) Natural IL-1Ra, predicted X1 and X2 mRNA (E), and protein (F) levels in overexpression KMM cells measured by RT-qPCR and ELISA, respectively. (G) Cell proliferation curves of KMM cells overexpressing IL-1Ra and variants described in D. (H and I) Analysis of cell cycle (H) and apoptosis (I) of KMM cells overexpressing IL-1Ra and variants described in D. Student’s t test was used for the analysis and results from 3 independent repeats are presented as mean ± s.d. (B, C, and E-I).

To determine whether dexamethasone induction of IL-1Ra was required for dexamethasone inhibition of cell proliferation of KSHV-transformed cells, we overexpressed IL-1Ra in KMM cells to mimic the function of dexamethasone. In humans, there are two IL-1Ra variants with one being secretory (sIL-1Ra) while the second one being intracellular (icIL-1Ra) (52). However, in rats, there are three IL-1Ra isoforms, including one natural IL-1Ra and two predicted isoforms (X1 and X2). Compared to the natural IL-1Ra transcript, the predicted isoforms X1 and X2 have alternative start codons, encoding proteins that have truncations of 19- or 35-amino acids in the N-terminals, respectively (Fig. 5D). Stable expression of individual IL-1Ra isoforms in KMM cells by lentiviral transduction (Fig. 5E and F) significantly inhibited the proliferation of KMM cells (Fig. 5G). However, these inhibitory effects were weak compared to dexamethasone treatment, ranging 13-28%. While these IL-1Ra isoforms appeared to inhibit cell cycle G0/G1 to S phase transition without inducing apoptotic cells, the effects were marginal without reaching the statistical significance (Fig. 5H and I). These results indicated that dexamethasone induction of IL-1Ra was unlikely the major reason for its inhibitory effect on the proliferation of KSHV-transformed cells.

### KSHV-encoded miRNAs mediate KSHV induction of IL-1α in KSHV-transformed cells

Our results showed that IL-1α was upregulated in KSHV-transformed cells and dexamethasone suppression of IL-1α was essential for its anti-proliferation function. Thus, we examined the viral gene(s) that might mediate KSHV induction of IL-1α. KSHV is in tight latency in KMM cells and only expresses a few latent genes including LANA (ORF73), vCyclin (ORF72), vFLIP (ORF71), and a miRNA cluster, all of which are essential for KSHV-induced cellular transformation (53-55). LANA is required for the persistence of viral episome and hence KSHV latent infection (56). Thus, genetic deletion of LANA in KSHV-infected cells is not feasible.

Deletion of the miRNA cluster (pre-miR-1-9 and -11) significantly reduced the level of IL-1α transcript, and overexpression of the miRNA cluster alone in MM cells was sufficient to significantly increase the IL-1α expression (Fig. 6A and B), indicating that the miRNA cluster contributed to KSHV induction of IL-1α. While deletion of vFLIP did not affect the expression of IL-1α transcript, overexpression of vFLIP in MM cells significantly increased the expression of IL-1α (Fig. 6A and C). These inconsistent results are likely due to the fact that vFLIP is usually expressed in extremely low level in KSHV latently infected cells, which can be overcome through overexpression (57). Therefore, although vFLIP was capable of inducing IL-1α, it unlikely contributed to KSHV induction of IL-1α. Deletion of vCyclin had a minor effect on IL-1α repression and overexpression of vCyclin in MM cells did not alter the level of IL-1α transcript (Fig. 6A and C), indicating that vCyclin unlikely contributed to KSHV induction of IL-1α.

**FIG 6.**
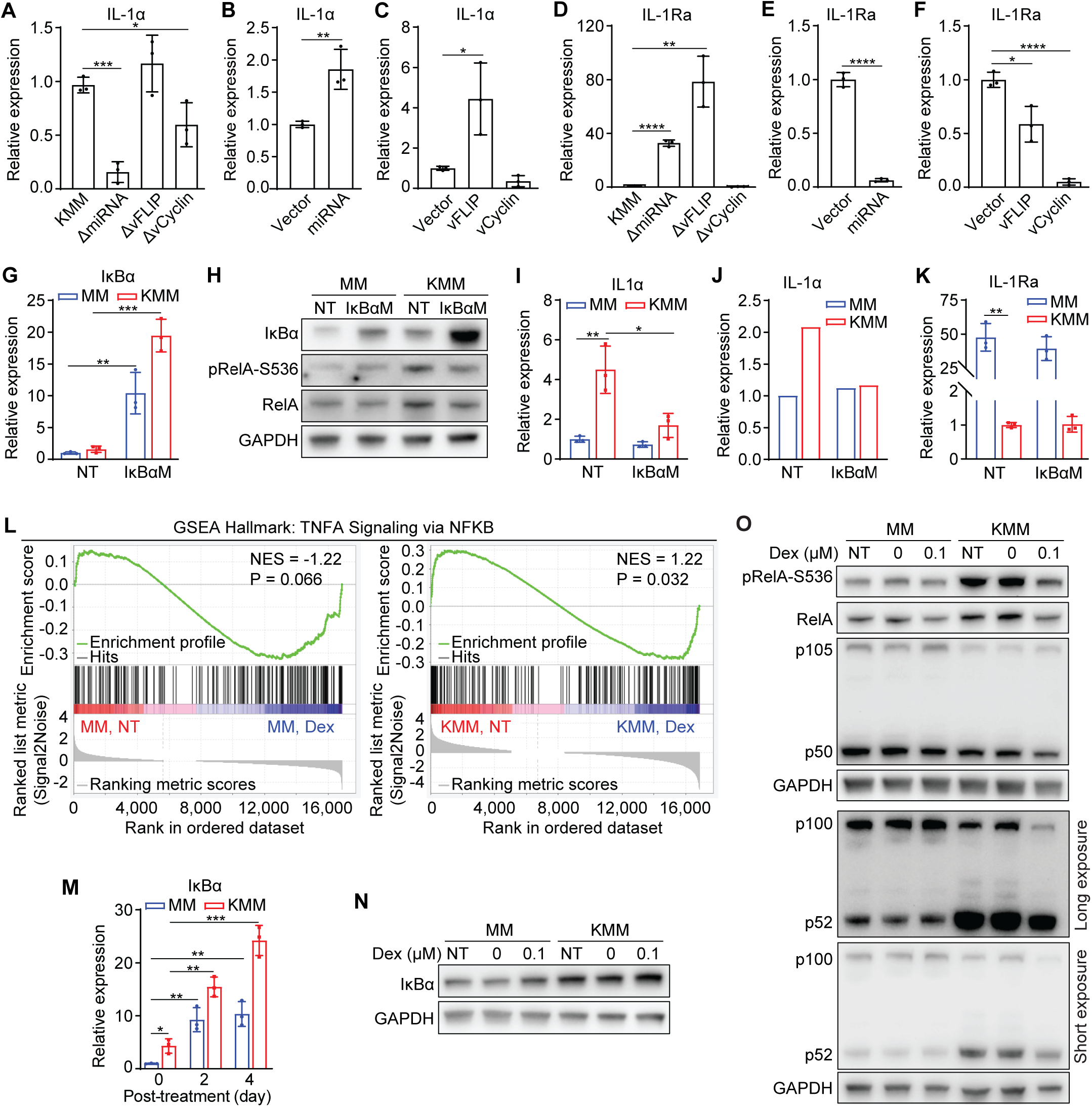
The NF-κB pathway mediates dexamethasone suppression of IL-1α induced by KSHV miRNAs and vFLIP. (A-C) IL-1α mRNA expression level measured by RT-qPCR in cells harboring wild-type KSHV (KMM), or KSHV with a deletion of the miRNA cluster (ΔmiRNA), vFLIP (ΔvFLIP) or vCyclin (ΔvCyclin) (A), MM cells expressing the miRNA cluster or vector control (B), or vFLIP, vCyclin or vector control (C). (D-F) IL-1Ra mRNA expression level measured by RT-qPCR in KMM cells, or cell harboring ΔmiRNA, ΔvFLIP or ΔvCyclin mutant (D), MM cells expressing the miRNA cluster or vector control (E), or vFLIP, vCyclin or Vector control (F). (G) IκBα mRNA expression levels in MM and KMM cells stably expressing IκBαM measured by RT-qPCR. (H) Immunoblots of IκBα, pRelA-S536 and RelA in MM and KMM cells stably expressing IκBαM. (I and J) IL-1α mRNA (I) and protein (J) levels in MM and KMM cells stably expressing IκBαM measured by RT-qPCR and ELISA, respectively. (K) IL-1Ra mRNA expression levels in MM and KMM cells stably expressing IκBαM measured by RT-qPCR. (L) Enrichment plots of TNFA signaling via NFKB (GSEA hallmark) of MM and KMM cells treated with dexamethasone. (M) IκBα mRNA expression levels in MM and KMM treated with 0 or 0.1 µM dexamethasone for 2 days measured by RT-qPCR. (N) Immunoblot of IκBα of MM and KMM cells treated with 0 or 0.1 µM dexamethasone for 2 days. (O) Immunoblots of pRelA-S536, RelA, p105/p50 and p100/p52 in MM and KMM cells treated with 0 or 0.1 µM dexamethasone for 2 days. Student’s t test was used for the analysis and results from 3 independent repeats are presented as mean ± s.d. (A-G, I, K and M).

Similarly, we examined viral genes that might mediate KSHV suppression of IL-1Ra. Deletion of either vFLIP or the miRNA cluster significantly increased the IL-1Ra transcripts while overexpression of either of them alone in MM cells significantly reduced the IL-1Ra expression (Fig. 6D-F), indicating that both vFLIP and the miRNA cluster were important for KSHV suppression of IL-1Ra. Interestingly, while deletion of vCyclin had no effect on IL-1Ra expression, overexpression of vCyclin in MM cells significantly reduced the IL-1Ra level (Fig. 6D-F). Similar to vFLIP, vCyclin is expressed at an extremely low level in KSHV latently infected cells (53). Thus, its suppressive effect on IL-1Ra expression only manifested following overexpression. Hence, while vCyclin was capable of suppressing IL-1Ra, it unlikely contributed to KSHV suppression of IL-1Ra.

### NF-κB activation mediates KSHV induction of IL-1α in KSHV-transformed cells

It has been reported that KSHV infection could activate the NF-κB pathway through both vFLIP and miRNAs (54, 58-61). Specifically, vFLIP regulates both classical and alternative NF-κB pathways by activating the IκB kinase (IKK) complex and promoting p100 processing into p52, respectively (58-60). KSHV miR-K1 and likely other viral miRNAs activate the NF-κB pathway by suppressing IκBα expression, an inhibitor of NF-κB complex (54, 61). Maximal KSHV activation of the NF-κB pathway requires both KSHV vFLIP and miRNAs (54). Thus, we examined the role of the NF-κB pathway in KSHV induction of IL-1α. We overexpressed a constitutively active form of IκBα with mutated phosphorylation sites (IκBαM), a super suppressor of the NF-κB pathway (62), in both MM and KMM cells (Fig. 6G and H). As expected, overexpression of IκBαM inhibited RelA phosphorylation at S536 (pRelA-S536) in KMM cells but had minimal effect on MM cells (Fig. 6H). Importantly, IκBαM suppression of the NF-κB pathway significantly decreased levels of IL-1α transcript and protein in KMM cells but had no effect on MM cells (Fig. 6I and J). Additionally, overexpression of IκBαM had no effect on IL-1Ra expression in both MM and KMM cells (Fig. 6K). These results indicated that the NF-κB pathway mediated KSHV induction of IL-1α but did not regulate KSHV suppression of IL-1Ra.

### Dexamethasone inhibits the NF-κB pathway by inducing IκBα in KSHV-transformed cells

Because KSHV latent genes are essential for KSHV-induced cell proliferation and cellular transformation, one possible mechanism of dexamethasone inhibition of cell proliferation is by suppressing the expression of KSHV latent genes. However, we did not observe any obvious expression changes of KSHV latent genes, including LANA, vCyclin and vFLIP as well as miR-K1, -K4, and -K11 in KMM cells following dexamethasone treatment (Fig. S2). There was also no expression change with the KSHV immediate-early lytic gene, the replication and transcription activator (RTA), polyadenylated nuclear (PAN) RNA, the early lytic gene ORF57, and the late lytic gene ORF65 (Fig. S2).

Because inhibition of the NF-κB pathway decreased the levels of IL-1α transcript and protein (Fig. 6I and J), we examined whether dexamethasone inhibited the NF-κB pathway to suppress IL-1α expression. The GSEA hallmark pathway analysis showed that TNF-α signaling via NF-κB was suppressed by dexamethasone (Fig. 6L). Examination of IκBα transcript and protein levels revealed that the IκBα expression was induced by dexamethasone in MM and KMM cells (Fig. 6M and N). These results were consistent with the reports that the *IKBA* gene encoding IκBα is a GR transcriptional target (63, 64). In fact, several GR-binding sites on the promoter of *IKBA* have been identified by ChIP-seq (65, 66). We further examined the NF-κB transcription factors, including RelA and p105/p50 of the classical NF-κB pathway, and p100/p52 of the alternative NF-κB pathway. As have been reported before, KSHV infection induced RelA expression and phosphorylation, and promoted p100-to-p52 processing to activate both classical and alternative NF-κB pathways (Fig. 6O). In agreement with the induction of IκBα, dexamethasone inhibited both classical and alternative NF-κB pathways (Fig. 6O). In addition, the p50 level was also reduced by dexamethasone in MM and KMM cells.

### KSHV genes regulate GR signaling in KSHV-transformed cells

While KSHV-transformed cells were responsive to dexamethasone, basal GR signaling was inhibited in cell cultures and in tumors (Fig. 1I and 2G). Specifically, we detected a lower level of pGR-S211 in KMM cells than MM cells (Fig. 1I), and almost no pGR-S211 signal in LANA-positive cells in KS-like tumors despite the presence of a robust signal in LANA-negative cells (Fig. 2G). Therefore, we examined viral genes that might mediate GR signaling in KSHV-transformed cells. Deletion of the miRNA cluster or vCyclin significantly reduced GR mRNA and protein levels (Fig. 7A and B). Interestingly, while deletion of vFLIP had no effect on GR mRNA, it increased GR protein level, suggesting the vFLIP might regulate GR expression at the post-transcriptional level. The results of pGR-S211 were consistent with those of total GR protein levels with deletion of the miRNA cluster or vCyclin predominant reducing GR phosphorylation while deletion of vFLIP enhancing GR signaling (Fig. 7B). We then determined whether expression of any single viral gene or the miRNA cluster in MM cells was sufficient to alter GR level and signaling. In agreement with the reverse genetic results, overexpression of the miRNA cluster increased while overexpression of vFLIP reduced the GR mRNA and protein levels as well as GR signaling (Fig. 7C-F). Interestingly, overexpression of vCyclin reduced the GR expression and phosphorylation, suggesting the distinct functions of vCyclin in the context of viral infection and single gene overexpression. These results suggested that vFLIP and possibly vCyclin likely contributed to KSHV inhibition of basal GR signaling in KSHV-transformed cells. However, the regulatory mechanisms of GR expression and signaling response were complex in the context of KSHV infection.

**FIG 7.**
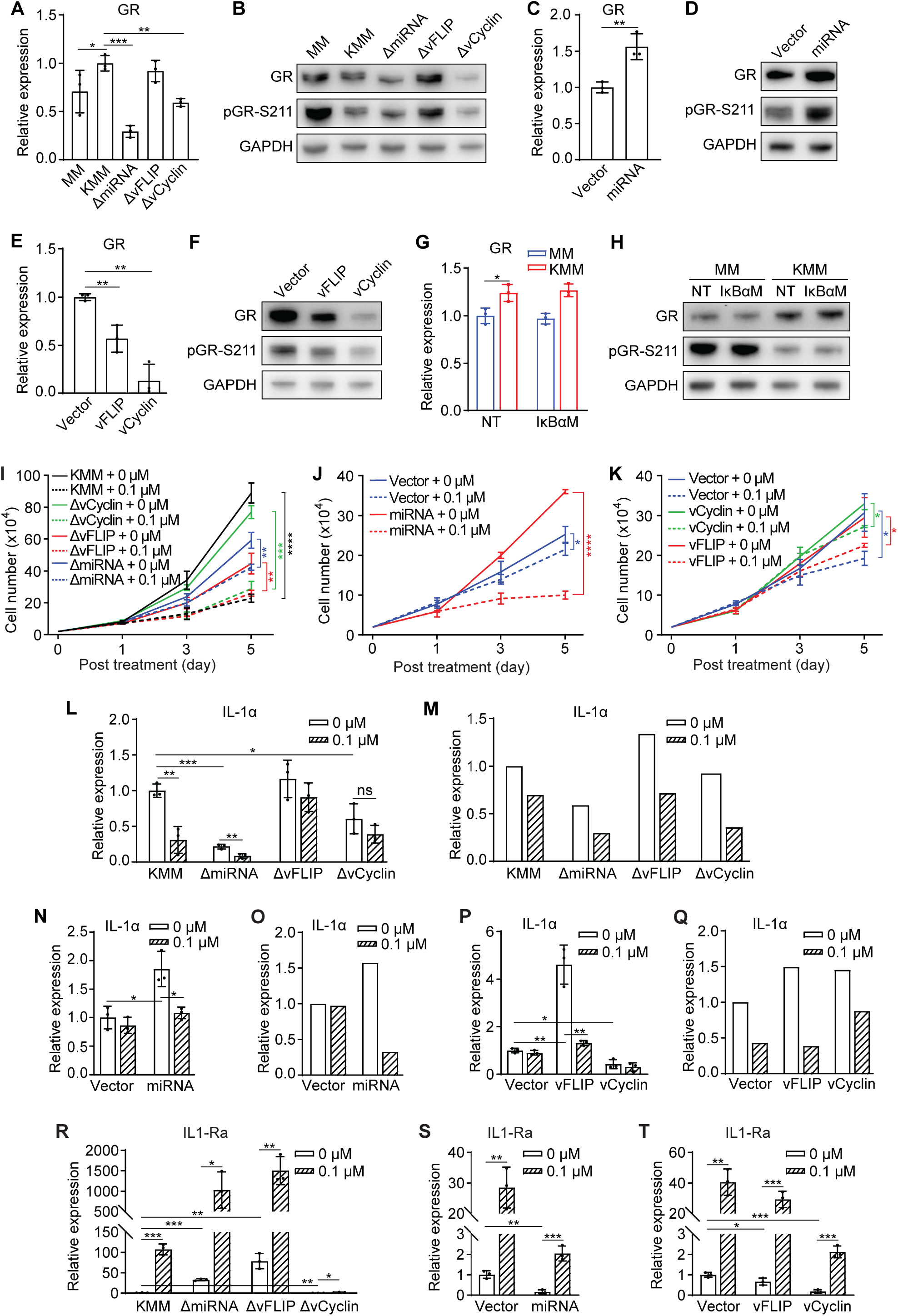
vFLIP inhibition of the GR pathway in KSHV-transformed cells does not impede cell responsiveness to dexamethasone. (A) GR mRNA expression levels in MM and KMM cells, or cells harboring ΔmiRNA, ΔvFLIP or ΔvCyclin mutant. (B) Immunoblots of GR and pGR-S211 in MM and KMM cells, or cells harboring ΔmiRNA, ΔvFLIP or ΔvCyclin mutant. (C) GR mRNA expression levels in MM cells expressing the miRNA cluster or the vector control. (D) Immunoblots of GR and pGR-S211 in MM cells expressing the miRNA cluster or the vector control. (E) GR mRNA expression levels in MM cells expressing vFLIP, vCyclin or the vector control. (F) Immunoblots of GR and pGR-S211 in MM cells expressing vFLIP, vCyclin or the vector control. (G) GR mRNA expression levels in MM and KMM cells stably expressing IκBαM or the vector control. (H) Immunoblots of GR and pGR-S211 in MM and KMM cells stably expressing IκBαM or the vector control. (I-K) Proliferation curves of KMM cells and cells harboring ΔmiRNA, ΔvFLIP or ΔvCyclin mutant (I), MM cells expressing the miRNA cluster or the vector control (J), or MM cells overexpressing vFLIP, vCyclin or the vector control (K) treated with 0 or 0.1 µM dexamethasone. (L and M) IL-1α mRNA (L) and protein (M) levels in KMM cells, and cells harboring ΔmiRNA, ΔvFLIP or ΔvCyclin mutant treated with 0 or 0.1 µM dexamethasone for 2 days measured by RT-qPCR and ELISA, respectively. (N and O) IL-1α mRNA (N) and protein (O) levels in MM cells expressing the miRNA cluster or vector control treated with 0 or 0.1 µM dexamethasone for 2 days measured by RT-qPCR and ELISA, respectively. (P and Q) IL-1α mRNA (P) and protein (Q) levels in MM cells expressing vFLIP, vCyclin or vector control treated with 0 or 0.1 µM dexamethasone for 2 days measured by RT-qPCR and ELISA, respectively. (R-T) IL-1Ra mRNA expression levels in KMM cells and cells harboring ΔmiRNA, ΔvFLIP or ΔvCyclin mutant (R), MM cells expressing the miRNA cluster or the vector control (S), or vFLIP, vCyclin or the vector control (T) treated with 0 or 0.1 µM dexamethasone for 2 days. Student’s t test was used for the analysis and results from 3 independent repeats are presented as mean ± s.d. (A, C, E, G, I-L, N, P and R-T).

Given that the NF-κB pathway mediated KSHV induction of IL-1α, we examined whether the NF-κB pathway might regulate GR expression and signaling. Suppression of the NF-κB complex by overexpressing IκBαM affected neither levels of GR transcript and protein nor GR signaling (Fig. 7G and H), indicating that the NF-κB pathway was not involved in regulating basal GR signaling in KSHV-transformed cells.

Because KSHV genes and miRNAs could regulate GR expression and signaling, we further examined whether they might mediate the cellular response to dexamethasone. Consistent with the results of our previous studies (53, 55), deletion of either the miRNA cluster or vFLIP compromised cell proliferation while deletion of vCyclin reduced cell proliferation at high cell density (Fig. 7I). However, cells harboring any of the mutant viruses remained responsive to the inhibitory effect of dexamethasone (Fig. 7I), again supporting the complex mechanism involved in regulating GR signaling in KSHV-transformed cells. In contrast, overexpression of the miRNA cluster in MM cells, which increased GR expression (Fig. 7C and D), significantly enhanced the responsiveness of cells to dexamethasone’s inhibitory effect from 14.85% to 72.22% (Fig. 7J). These results were consistent with the observations that the KSHV miRNAs are proinflammatory (67-69). Nevertheless, overexpression of vFLIP or vCyclin did not alter the cellular responsiveness to dexamethasone treatment (Fig. 7K), which were consistent with the attenuation of GR signaling by these two viral genes (Fig. 7F).

Finally, we determined whether KSHV genes might regulate the inhibitory effect of dexamethasone on IL-1α expression. Dexamethasone treatment reduced IL-1α transcript and protein in cells harboring any of the mutant viruses (Fig. 7L and M), which was in agreement with the results of cell proliferation (Fig. 7I). Furthermore, overexpression of the miRNA cluster, vFLIP or vCyclin did not alter dexamethasone’s suppression of IL-1α expression at both transcript and protein levels (Fig. 7N-Q). Meanwhile, dexamethasone treatment increased IL-1Ra expression in cells harboring the miRNA and vFLIP mutants but not vCyclin mutant (Fig. 7R), whereas cells with overexpression of the miRNA cluster, vFLIP or vCyclin remained responsive to dexamethasone induction of IL-1Ra (Fig. 7S and T). Taken together, vFLIP, vCyclin and the miRNA cluster did not regulate the responsiveness to dexamethasone treatment, which was likely due to the intrinsic inflammatory properties of KSHV-transformed cells.

## Discussion

Although hyperinflammation is recognized as a pathological hallmark in KS tumors, how it is induced remains unclear. HIV infection is known to cause immunosuppression, induce metabolic disorders, and enhance viral and bacterial coinfections, all of which could contribute to inflammation in AIDS-KS tumors (16, 17, 70). KSHV primary infection, expression of numerous KSHV genes, and abnormal activation of immune response in KSHV-infected cells including the alternative complement system, and toll-like receptor 4 and 5 (TLR4 and TLR5) pathways could also trigger inflammation (21, 28-31, 38-40). Nevertheless, KSHV induction of inflammation has never been examined in a KSHV-induced cellular transformation system. In this study, using a model of KSHV-induced cellular transformation of primary cells (41), we have shown the activation of numerous inflammatory pathways in KSHV-transformed cells (Fig. 1B and C). Among them, the IL-1 signaling pathway has been implicated in many disease conditions (71, 72). We have identified IL-1α as the most highly upregulated gene while IL-1Ra as the most highly downregulated gene in this pathway in KSHV-transformed cells (Fig. 4A-C, 5A and B).

Because inflammation might promote KS development by enhancing both KSHV lytic replication at the early stage of KS and the proliferation of latently infected KS tumor cells (17, 37-40), we have examined the role of inflammation in KSHV-induced cellular transformation using the anti-inflammatory agent dexamethasone. We found that dexamethasone inhibited the proliferation and colony formation in soft agar of KSHV-transformed cells (Fig. 1D and E). Consistently, dexamethasone induced cell cycle arrest and inhibited the AKT/mTORC1 pathway (Fig. 1F and H). *In vivo*, dexamethasone suppressed the initiation and growth of KSHV-induced tumors (Fig. 2B-E). Importantly, dexamethasone inhibited KSHV-induced inflammation by suppressing IL-1α and inducing IL-1Ra (Fig. 4A-C, 5A and B). Indeed, treatment with recombinant IL-1α protein was sufficient to enhance cell proliferation in both MM and KMM cells (Fig. 4F and G).

Compared to the IL-1R ligand IL-1β, less is known on the role of IL-1α-dependent proinflammation in cancer despite it has become an apical regulator of inflammatory response in a much wider cellular scenario due to its unique properties (71-73). To our knowledge, this is the first study to delineate the mechanism of IL-1α induction and functionally examine its role in KSHV-induced cellular transformation. IL-1α can be rapidly induced by a wide variety of stimulation including pathogen infections. Interestingly, unlike IL-1β, which must be processed to become mature for release, both precursor and mature IL-1α are bioactive and function as secreted or membrane bound cytokines (74-76). Furthermore, IL-1α can also be translocated to nucleus to modulate gene expression in an IL-1R-independent manner (77, 78). Despite the important role of IL-1α in human diseases, the mechanisms regulating IL-1α synthesis and secretion remain unclear. Given its important role in different conditions, understanding the biology of IL-1α could help develop novel therapies, particularly, for inflammatory diseases.

By using reverse genetics, we have shown that KSHV miRNAs are necessary and sufficient for KSHV induction of IL-1α, most likely through the NF-κB pathway. Meanwhile, KSHV miRNAs and vFLIP are necessary and sufficient for KSHV suppression of IL-1Ra. Given the anti-inflammation role of GR signaling, we have examined the status of the GR pathway in KSHV-transformed cells. We have found that the GR pathway is suppressed in KSHV-transformed cells (Fig. 1I, 2G and 7B). KSHV vFLIP and likely vCyclin mediate KSHV suppression of GR signaling through an unknown mechanism (Fig. 7A, B, E and F). Thus, both KSHV induction of inflammation and suppression of anti-inflammation contribute to the hyperinflammation in KSHV-transformed cells (Fig. 8).

**FIG 8.**
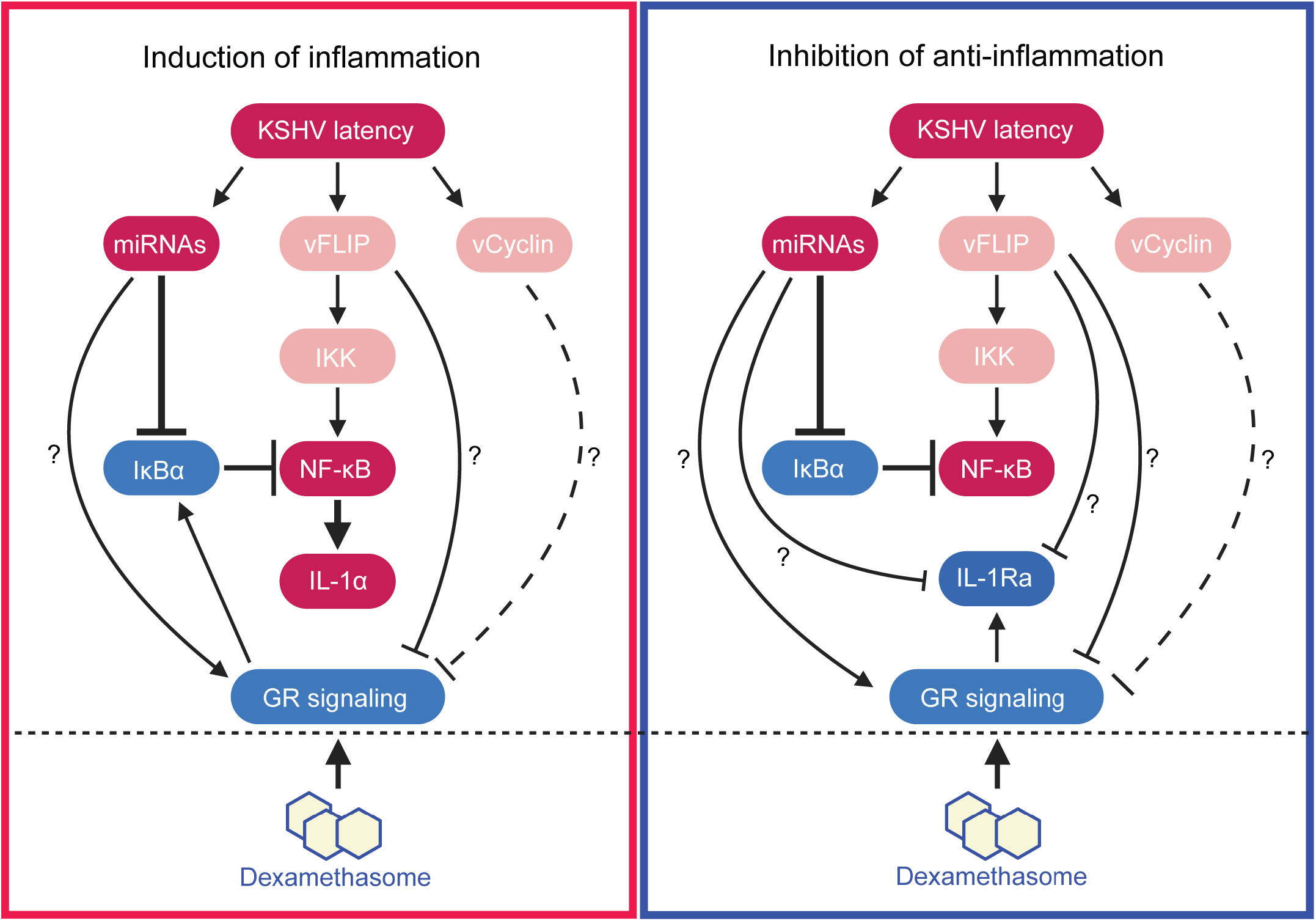
A working model illustrating the mechanisms mediating KSHV induction of hyperinflammation, and dexamethasone suppression of KSHV-induced inflammation and cellular transformation.

Meanwhile, dexamethasone also inhibits KSHV-activated NF-κB pathways through GR-transactivation of its suppressor IκBα (Fig. 6H-J). The NF-κB pathway is essential for the survival of KSHV-transformed cells (54, 55). As expected, a functional GR signaling pathway is required for the action of dexamethasone (Fig. 1K, 4D, 4E and 5C). However, GR knockdown did not affect the basal expression of IL-1α and IL-1Ra in KMM cells, which could be due to the fact that GR signaling was already inhibited in these transformed cells (Fig. 1I, 2G and 7B). While the GR level in KMM cells was reduced compared to MM cells, KMM cells remained responsive to GR ligands (Fig. 1I). Upon dexamethasone treatment, GR was activated and rapidly translocated into the nucleus. Activated GR binds to specific glucocorticoid response elements (GREs) in the promoter region of target genes (43). Both IL-1Ra and IκBα genes were induced by dexamethasone (Fig. 5A and B, 6M and N). These results are consistent with the presence of GREs in the promoters of IL-1Ra and IκBα genes identified by ChIP-seq (65, 66). Dexamethasone induction of IκBα gene inhibited the NF-κB pathway resulting in the suppression of IL-1α (Fig. 6H-J, M-O). Thus, the interaction of GR and NF-κB pathways mediate dexamethasone’s anti-inflammatory and anti-cellular transformation functions in KSHV-transformed cells by directly inducing the expression of IL-1Ra but indirectly inhibiting the induction of IL-1α (Fig. 8). Because both GR and NF-κB pathways are key mediators of inflammatory response, it is not surprising that the two pathways intersect to mediate KSHV-induced inflammation and cellular transformation. On the other hand, there was no evidence that the NF-κB pathway directly affected GR expression and signaling (Fig. 7G and H).

In summary, we have shown the activation of numerous inflammatory pathways and suppression of the anti-inflammatory GR pathway in KSHV-transformed cells, and that induction IL-1α and suppression of IL-1Ra in the IL-1 pathway are essential for KSHV-induced cellular transformation and tumorigenesis (Fig. 8). Mechanistically, we have shown that the NF-κB pathway mediates KSHV induction of IL-1α while the GR pathway mediates KSHV suppression of IL-1Ra. Consequently, anti-inflammatory agents such as dexamethasone is effective for inhibiting KSHV-induced inflammation and cellular transformation, at least in part by suppressing IL-1α induction and releasing IL-1Ra suppression (Fig. 8). Our results illustrate inflammation as a viable therapeutic target for preventing and treating KSHV-induced malignancies. This approach is particularly valuable for populations with economic disadvantage and poor access to health care since many anti-inflammatory agents such as dexamethasone are widely available and affordable.

## Materials and Methods

### Cell lines and cell culture

Rat primary embryonic metanephric mesenchymal precursor cells (MM) and KSHV-transformed MM cells (KMM) (41), MM cells infected by KSHV mutant with deletion of the miRNA cluster (ΔmiRNA) (54), vFLIP (ΔvFLIP) (57) or vCyclin (ΔvCyclin) deletion (53) were used in this study. MM and 293T cells were cultured in Dulbecco’s Modified Eagle Medium (DMEM) with 10% fetal bovine serum (Sigma-Aldrich, F2442) and 1% Penicillin-Streptomycin solution. KMM cells and cells harboring different KSHV mutants were cultured in the same medium with 250 µg/mL hygromycin. Cells were treated with ethanol, dexamethasone (Sigma-Aldrich, D1756) or recombinant IL-1α protein (Novus Biologicals, NBP2-35227) as described.

### Soft agar assay

The soft agar assay was performed as previously described (41). Agar was prepared with culture medium containing 20% FBS. A total of 1.5 mL of 0.5% agar was plated into each well of 6-well plates to form the bottom layer, then covered with 2 mL of 0.3% top layer agar containing 2 × 10^4^ cells. Dexamethasone or ethanol was added to the medium. At day 15, the plates were photographed with an Olympus inverted microscope using a 2x objective.

### Apoptosis and cell cycle assays

Apoptotic cells were detected by flow cytometry using Fixable Viability Dye eFluor™ 660 (Invitrogen, 65-0864) and Annexin V Apoptosis Detection Kits (Invitrogen, 88-8103-74). Cell cycle was examined by flow cytometry following 5′-bromo-2′-deoxyuridine (BrdU) labeling (BrdU: Sigma-Aldrich, MB5002; BrdU antibody: Invitrogen, B35129) and propidium iodide (PI) staining (Sigma-Aldrich, P4864). The data were analyzed using the FlowJo software.

### RNA sequencing (RNA-seq) and data processing

Total RNA was extracted from cells using TRI reagent (Sigma-Aldrich, T9424) according to the manufacturer’s instructions. mRNA was isolated using Oligo(dT) beads and library was prepared following the Illumina TruSeq Stranded mRNA Library Prep Kit (Illumina, 20020594). Sequencing was carried out using Illumina HiSeq 3000 with the 50-bp single-read sequencing module. NCBI genome assembly mRatBN7.2 was used as a reference genome. The gene expression analysis was processed using the QIAGEN CLC Workbench. Transcripts per million (TPM) was used for normalizing sample dependencies, and calculating the differential expression levels. The pathway analysis was conducted using DAVID Bioinformatics Resources and Gene Set Enrichment Analysis (GSEA).

### Immunoblotting

Cell pellets were lysed in 1x Laemmli buffer and protein lysates were resolved in SDS polyacrylamide gels and transferred to nitrocellulose membranes (Cytiva, 10600004). Antibodies used for these experiments were as follows: GAPDH (CST, 5174), β-tubulin (Sigma, 7B9), p27 (CST, 3686), cyclin D1 (CST, 2978), AKT (CST, 4691), pAKT-T308 (CST, 2965), GR (CST, 12041), pGR-S211 (CST, 4161), GILZ (Invitrogen, PA5-93215), p105/p50 (CST, 13586), p100/p52 (CST, 4882), IκBα (CST, 4814), RelA (CST, 8242), and pRelA-S536 (CST, 3033).

### Reverse transcription real-time quantitative PCR (RT-qPCR)

Total RNA was extracted from cells using the TRI reagent (Sigma-Aldrich, T9424) according to the manufacturer’s instructions. The reverse transcription was carried out with the cDNA Synthesis Kit (Thermo Scientific, K1652). The Universal SYBR Green Supermix (Bio-Rad, 172-5272) was used for real-time quantitative PCR (qPCR). The primers were as follows (for Rattus norvegicus): β-actin (F, 5’-CCATGTACCCAGGCATTGCT-3’; R, 5’-AGCCACCAATCCACACAGAG-3’), IL-1α (F, 5’-TCAAGATGGCCAAAGTTCCTGA-3’; R, 5’-AGACAGATGGTCAATGGCAGA-3’), IL-1Ra (F, 5’-GAATGTGTTCTTGGGCATCC-3’; R, 5’-TGTTGTGCAGAGGAACCATC-3’), IκBα (F, 5’-TGGCCAGTGTAGCAGTCTTG-3’; R, 5’-GTGTGGCCGTTGTAGTTGGT-3’), and GR (F, 5’-ACTCAAGCCCTGGAATGAGAC-3’; R, 5’-GCTGGGCAGTTTTTCCTTCG-3’).

### siRNA-mediated GR knockdown

The small interfering RNAs (siRNAs) against Rattus norvegicus GR (SASI_Rn01_00092098, SASI_Rn01_00092099) and scramble control (SIC001) were obtained from Sigma-Aldrich. Lipofectamine 2000 Kit (Invitrogen, 11668019) was used for the delivery of siRNAs according to the manufacturer’s instructions. Knockdown efficiency was examined at 48 h post transfection.

### IL-1Ra and IκBαM overexpression

Rat IL-1Ra isoforms were cloned into pCDH vector between EcoRI and BamHI sites with a 3x FLAG tag at the N-terminus. The construct of the NF-κB super suppressor IκBαM containing a N-terminal S36 mutation and a C-terminal PEST mutation was a gift from Dr. Inder Verma (62). Lentivirus was produced by transfecting the expression plasmid with p8.74/pMDG lentiviral packaging system into 293T cells using the Lipofectamine 2000 Kit (Invitrogen, 11668019). Supernatant containing the lentivirus was harvested at 48 and 72 h post-transduction, and filtered through a 0.45 µM filter. Lentiviral transduction was carried out at a MOI of 6 in the presence of 10 μg/ml polybrene by spinning infection at 1,500 rpm for 1 h. The expression efficiency was examined at 72 h post transduction. Puromycin at 2.5 µg/mL was added to the culture medium for selecting stable cell cultures.

### Enzyme-linked immunosorbent assay (ELISA)

Cell culture supernatants were collected and centrifuged at 1,500 rpm for 15 min at 4 °C. Cell pellets were lysed in extraction buffer (100 mM Tris pH 7.4, 150 mM NaCl, 1 mM EGTA, 1 mM EDTA, 1% Triton X-100, 0.5% sodium deoxycholate, and a cocktail of phosphatase and protease inhibitors). ELISA was performed using the rat IL-1α ELISA Kit (Invitrogen, BMS627) and IL-1Ra ELISA Kit (Invitrogen, ERA22RB) according to the manufacturer’s instructions. The results were normalized by total protein concentrations of individual samples.

### Animal experiments

Female nude mice at 4-5 weeks old were purchased from Envigo. KMM cells were injected into both flanks of 32 mice at 10^7^ cells per site. Each mouse generates two tumors. In Stage I experiment, mice were randomly split into two groups to receive either 20 mg/kg dexamethasone (Dex, n=16) or vehicle treatment (ethanol in phosphate buffer saline, Vehicle, n=16) starting at day 17 post-inoculation by intravenous injection (IP) for 5 days per week (day 17 to 50). In Stage II experiment, the vehicle group was evenly divided into two subgroups at day 56, and one subgroup was treated with dexamethasone (Vehicle-Dex, n=8) while the second subgroup remained untreated (Vehicle-Vehicle, n=8). Meanwhile, the dexamethasone group was evenly divided into two subgroups, and one subgroup remained treated with dexamethasone (Dex-Dex, n=8) while the second subgroup was no longer treated (Dex-Vehicle, n=8). The initial vehicle group were terminated at day 77 while the initial dexamethasone group were terminated at day 140, when the volumes of some tumors reached 1.5 cm^3^. Tumor incidence and growth were analyzed as previously described (41). The body weights of the mice and tumor volumes were measured every three days.

### Immunofluorescence (IF) and immunohistochemistry (IHC) staining

IF and IHC were carried out as previously described (79). Antibodies used for these experiments were as follows: Ki-67 (Abcam, 16667), GR (CST, 12041), pGR-S211 (CST, 4161), LANA (Abcam, 4103) and IL-1α (LSBio, LS-C177212). DAPI (Sigma-Aldrich, D9542) was used for nuclear counterstaining.

### Statistical analysis

Results were presented as the mean ± standard error of the mean. The differences between two groups were analyzed using Student’s t-test, and one-way analysis of variance was performed when more than two groups were compared. Statistical tests were two-sided. A P < 0.05 was considered statistically significant. Statistical symbols “*,” “**,” “***” and “****” represent P < 0.05, < 0.01, < 0.001 and < 0.0001, respectively, while “NS” indicates “not significant.”

## SUPPLEMENTAL MATERIAL

Supplemental material is available online only.

FIG S1, PDF file, 128 KB.

FIG S2, PDF file, 170 KB.

Table S1, Excel file, 450 KB.

Table S2, Excel file, 209 KB.

Table S3, Excel file, 312 KB.

Table S4, Excel file, 274 KB.

Table S5, Excel file, 47 KB.

Table S6, Excel file, 81 KB.

## Acknowledgments

We would like to thank Dr. Inder Verma for the super suppressor IκBαM construct, and members from Dr. Gao’s laboratory for technical supports and helpful suggestions. This work was supported by grants from National Institutes of Health (CA096512, CA278812, CA284554 and CA124332 to S.-J. Gao), and in part by award P30CA047904.

## Authors’ Contributions

L. P. Chen: Investigation, methodology, data curation, formal analysis, visualization, writing–original draft, writing–review. L. Ding: Investigation, data curation, writing–review. X. Wang: Investigation, data curation, writing–review. Y. F. Huang: Formal analysis, writing–review and editing. S.-J. Gao: Conceptualization, managing, formal analysis, investigation, visualization, methodology, writing–original draft, writing– review and editing.

## SUPPLEMENTAL MATERIAL

### Supplemental Tables

**Table S1** Differentially expressed genes in KMM cells compared to MM cells

**Table S2** Genes in enriched pathways in KMM cells compared to MM cells analyzed using GOBP database

**Table S3** Differentially expressed genes in MM cells treated with dexamethasone

**Table S4** Differentially expressed genes in KMM cells treated with dexamethasone

**Table S5** Genes in the enriched pathways in MM cells treated dexamethasone analyzed using GOBP database

**Table S6** Genes in the enriched pathways in KMM cells treated dexamethasone analyzed using GOBP database

### Supplemental Figure Legends

**FIG S1** Body weights of mice treated with vehicle and dexamethasone (Dex) described in Fig 2. (A) Body weights of mice in Stage I experiment. (B) Body weights of mice in Stage II tumor regression experiment. (C) Body weights of mice in Stage II maintenance experiment.

**FIG S2** Expression levels of KSHV genes treated with dexamethasone. (A) LANA. (B) vCyclin. (C) vFLIP. (D) RTA. (E) PAN RNA. (F) ORF57. (G) ORF65. (H) miR-K1. (I) miR-K4. (J) miR-K11.

